# Microbial composition in larval water enhances *Aedes aegypti* development but reduces transmissibility of Zika virus

**DOI:** 10.1101/2021.08.03.455011

**Authors:** William Louie, Lark L. Coffey

**Affiliations:** University of California, Davis, School of Veterinary Medicine, Department of Pathology, Microbiology, and Immunology

## Abstract

Arthropod-borne viral (arboviral) pathogens comprise a significant global disease burden, and outbreaks are expected to increase as vectors expand. Surveillance and mitigation of arboviruses like Zika virus (ZIKV) require accurate estimates of transmissibility by vector mosquitoes. While numerous laboratory vector competence experiments show that *Aedes spp*. mosquitoes are competent ZIKV vectors, differences in experimental protocols prevent direct comparisons of relative transmissibility across studies. An understudied factor complicating these comparisons is differential environmental microbiota exposures, where most vector competence studies use mosquitoes reared in laboratory tap water, which does not represent the microbial complexity of environmental water where wild larvae develop. We simulated natural larval development by rearing Californian *Aedes aegypti* with microbes obtained from cemetery headstone water, a common larval habitat in California, compared to conventional laboratory tap water. *Ae. aegypti* larvae reared in environmental cemetery water pupated 3 days faster and at higher rates. Female adult mosquitoes reared in environmental water were less competent vectors of ZIKV compared to laboratory water-reared *Ae. aegypti*, as evidenced by significantly reduced infection and transmission rates for two 2015 ZIKV strains and in two *Ae. aegypti* colonies from California. Microbiome comparisons of laboratory- and environment-water reared mosquitoes as well as their rearing water showed significantly higher bacterial diversity in environment water; despite this pattern, corresponding differences in diversity were not consistently detected in adult mosquitoes reared in different water sources. We also detected more significant associations between the microbial composition of adult mosquitoes and whether they ingested a bloodmeal than larval water type. Together, these results highlight the role of transient microbes in the larval environment in modulating vector competence. Laboratory vector competence likely overestimates true transmissibility of arboviruses like ZIKV when conventional laboratory water is used for rearing.

**Importance:** We observed that *Ae. aegypti* mosquitoes reared in water from cemetery headstones instead of the laboratory tap exhibited a reduced capacity to become infected with and transmit Zika virus. Water from the environment contained more bacterial species than tap water, but these bacteria were not consistently detected in adult mosquitoes. Our results suggest that rearing mosquito larvae in water collected from local environments as opposed to laboratory tap water, as is conventional, provides a more realistic assessment of vector competence since it better recapitulates the natural environment in which larvae develop. Given that laboratory vector competence is used to define the species to target for control, use of environmental water to rear larvae could better approximate the microbial exposures of wild mosquitoes, lessening the potential for overestimating transmission risk.

## Introduction

Global expansion of arthropod borne viruses (arboviruses) poses a significant public health threat. Climate change and rapid urbanization may accelerate zoonotic spillover or re-emergence of arboviruses, increasing outbreaks in humans (1–3). Zika virus (*Flaviviridae, Flavivirus*, ZIKV) which was understudied since its discovery in 1947 in Uganda (4), garnered worldwide attention following outbreaks in 2015-16 (3, 5, 6). The wave of ZIKV epidemics, accompanied by newly recognized teratogenic phenotypes wherein ZIKV causes adverse outcomes in fetuses from infected pregnant mothers, now referred to as Congenital Zika Syndrome (7, 8), fueled efforts to better understand and mitigate transmission to curtail disease. Although the ZIKV pandemic of 2015-16 has ended, ZIKV may re-emerge via increased numbers of immunologically naïve people and geographic expansion of *Aedes spp*. vectors (9, 10).

Determining the ability of a mosquito to become infected by and transmit a virus (vector competence), is crucial for guiding surveillance and control, including to identify mosquito species to monitor and eliminate, and to model outbreak risk. Evaluating vector competence in the laboratory entails exposing mosquitoes to an infectious bloodmeal, followed by detection of viral RNA or infectious virus in mosquito tissues and saliva after an incubation period of 3-14 days. Mosquito-borne arboviruses must escape the mosquito midgut, infect the salivary glands, and be secreted into saliva for transmission. Although the approach to assessing laboratory vector competence is standard, outcomes across studies vary greatly (6), and may be influenced by virus strain and passage history (11), virus dose (12), mosquito species (13), intraspecies mosquito genetics (14, 15), larval nutrition and competition (16, 17), and incubation temperature (18–20). Since 2017, many vector competence studies have been performed using *Ae. aegypti* and *Ae. albopictus* from varied geographic origins and post-2015 strains of ZIKV (21–28). The absence of uniformity in the variables involved in laboratory vector competence makes direct comparisons across studies difficult. However, such comparisons are needed to assess reproducibility and to identify differences in vector competence across geographies.

The mosquito microbiome is an important variable that influences arbovirus vector competence, wherein specific taxa can modify it. *Ae. aegypti* infected with the bacterial endosymbiont *Wolbachia* fail to transmit ZIKV, dengue (DENV), and chikungunya virus (CHIKV), prompting field trials and experimental release of *Wolbachia-infected* male mosquitoes as a means of population replacement (29). Similarly, the bacterium *Chromobacterium Csp_P* reduces transmission of DENV by *Ae. aegypti*, and *Plasmodium falciparum*, the cause of malaria, by *Anopheles gambiae* (30). Members of the bacterial genus *Asaia* may also confer resistance of mosquitoes to arboviruses and *Plasmodium* (31–33). *Ae. aegypti* colonized with *Serratia* bacteria are more susceptible to infection by DENV and CHIKV *in vivo*, but less susceptible to ZIKV *in vitro* (34–37). However, the functional roles of specific microbial strains in modulating vector competence of mosquitoes in nature, where gut microbes exist as a community rather than as a monoculture, remains unclear. Examination of microbial strains in gnotobiotic mosquitoes requires repeatability in a microbial community context, including in the aqueous larval form. To address this gap, we analyzed microbial structures of larval *Ae. aegypti* to elucidate community dynamics of microbes that colonize larvae and adults, and then we assessed how differences in larval rearing environments and microbial composition affect ZIKV vector competence.

We modified the *Ae. aegypti* larval rearing environment by introducing microbes at different diversity and abundance. Since microbes in mosquitoes are primarily acquired through the environment (38, 39), rearing *Ae. aegypti* in different water sources provides control of microbial input to *Ae. aegypti* colonies in the laboratory (40, 41). Previous work showed that the bacterial microbiota of field-caught *Aedes* varies geographically (42) and that rearing field mosquitoes in a laboratory setting results in convergence of gut microbiota in just one generation (43). Moreover, larvae-acquired microbes play a significant role in larval development, where axenically (raised as a single organism, free of any microbes) reared mosquitoes exhibit inconsistent pupation success and reduced adult size, likely due to lack of nutritional supplementation by larval gut microbes (38, 44). Additionally, some larval gut microbes are passed transstadially to adults, suggesting that some microbes are symbiotic with mosquitoes through multiple life stages (45, 46). Consequently, microbes acquired by larvae are expected to influence *Aedes* mosquito physiology and immune status (47–49) which, along with direct physical interactions by microbes, is expected to impact ZIKV vector competence (50, 51). We used larval rearing water that we determined contained relatively low microbial content compared to microbe-rich water collected from outdoor environments in which *Ae. aegypti* larvae are naturally found to determine whether differences in water source influence ZIKV vector competence in a controlled mosquito genetic background. Our data show that reduced microbial exposure in colonized mosquitoes reared in laboratory water modulates vector competence and could explain variability in vector competence between laboratory and field mosquitoes.

## Materials and Methods

### Biosafety

All ZIKV experiments were conducted in a biosafety level 3 laboratory and were approved by the University of California, Davis, under Biological Use Authorization #R1863.

### Mosquitoes

Two sources of *Ae. aegypti* mosquitoes were used in this study. *Ae. aegypti* were field-collected as larvae in Los Angeles, CA, or as eggs in Clovis, CA, in 2016 and reared under standard insectary conditions for several generations until F_13-16_ and F_9_ eggs, respectively, were collected for use. Adults were morphologically identified by personnel training in recognizing *Ae. aegypti*. Insectary conditions during the laboratory colonization process were 26°C, 80% humidity, and 12 hour:12 hour light:dark cycle, with larvae maintained in 1 liter (L) deionized water (diH_2_O) at 200-400 larvae per pan and provided 1 pinch of fish food (Tetra, Melle, Germany) every other day until pupation. Adults were maintained in 30 x 30 x 30 cm mesh cages (BugDorm, Megaview Science, Taiwan) with constant access to 10% sucrose, all under septic conditions.

### Mosquito rearing

Urban-adapted *Ae. aegypti* larvae are known to develop within open containers including cemetery headstones, plant pots, rain barrels, abandoned tires, and bromeliads, which tend to accumulate nutrients and organic matter (17, 52–54). Outdoor and laboratory water sources were used in this study (Supplemental Table 1). For the laboratory water, ethanol-cleaned plastic trays were filled with 1 L laboratory tap deionized water (diH_2_O) in an insectary. Environmental water consisted of 2-3 L per collection of stagnant water from headstone receptacles in Davis Cemetery (Davis, CA) after rainfall. Separate water collections were conducted prior to each experiment to encompass variation in outdoor environmental conditions over time. The cemetery water was filtered through 1 mm mesh to remove insects, larvae, and large particulates and then centrifuged at 3000 g for 30 minutes to pellet microbes. The supernatant was discarded, and pellets were washed with sterile 1X phosphate buffered saline (PBS, Thermo Fisher Scientific, Emeryville, CA) three times prior to creating glycerol stocks that were frozen for later use. Environmental water aliquots were also plated on LB agar plates (Sigma-Aldrich, St. Louis, MO) to estimate live bacteria quantities prior to freezing at −80°C.

Mosquito eggs were surface sterilized by submerging in 5% bleach (Clorox, Oakland, CA) for 10 minutes, then washed twice in 70% ethanol (Thermo Fisher Scientific, Emeryville, CA) and dried for 10 minutes before hatching in diH_2_O. A PBS wash on a subset of eggs after surface sterilization was cultured on LB to confirm the removal of live bacteria from egg surfaces. Hatching was stimulated by either a pinch of active dry yeast (Red Star Yeast, Milwaukee, WI) in larval water or by inducing negative pressure (Rocker 400 Vacuum Pump, Sterlitech Corp., Kent, WA) to reduce the dissolved oxygen content for 30 minutes. A total of ~2500 larvae were transferred to six 1 L pans to achieve a density of 400-500 larvae/pan. Food was prepared in agarose plugs that were made by mixing 1% agarose (Sigma-Aldrich, St. Louis, MO) with pulverized fish food (final concentration of 100 g/L, or 10% [Tetra, Melle, Germany]) and rodent chow (final concentration of 80 g/L, or 8% [Teklad Global 18% Protein Rodent Diet, Envigo, Indianapolis, IN]), which was then autoclave-sterilized before casting into 12-well plates, a modification of a described approach (44) for standardizing larval diet. One plug was fed to larvae in each pan every other day. Pupae were counted once daily and transferred into plastic dishes containing sterile diH_2_O within 30 cm^2^ cloth cages. Once cages reached a mosquito density of about 500, adult females were transferred in batches of 100 to 32-ounce plastic containers (Amazon, Seattle, WA) for vector competence experiments. Larval trays and adult mosquitoes were maintained at 26°C, 80% humidity, and 12 hour:12 hour light:dark cycle for the duration of the experiment. All trays and adult mosquitoes were housed in the same incubator. Adult mosquitoes were provided constant access to filter-sterilized 10% sucrose (Thermo Fisher Scientific, Emeryville, CA).

### Virus sources and titrations

Two Asian-lineage ZIKV strains were used: PR15 (Puerto Rico 2015, PRVABC59, GenBank Accession # KX601168) and BR15 (Brazil 2015, SPH2015, GenBank Accession # KU321639), both of which were isolated from human serum and passaged 3 times in Vero cells (CCL-81, ATCC, Manassas, VA) before freezing in stocks. Stocks were titrated on Vero cells prior to bloodmeal presentation to confirm titers. Remaining bloodmeals were recovered after presentation to mosquitoes, frozen at −80°C, and back-titrated by plaque assay on Vero cells to confirm the administered dose. For titrations, bloodmeals were serially diluted 10-fold in DMEM, inoculated into one well, and incubated for 1 hour at 37°C in 5% CO_2_ with rocking every 15 min to prevent cell death due to desiccation. After 1 hour, 3 mL of 0.5% agarose (Thermo Fisher Scientific, Emeryville, CA) mixed with DMEM supplemented with 2% FBS and penicillin/streptomycin (Thermo Fisher Scientific, Emeryville, CA) was added to each well to generate a solid agar plug. The cells were incubated for 7 days at 37°C in 5% CO2, after which they were fixed with 4% formalin (Thermo Fisher Scientific, Emeryville, CA) for 30 minutes, plugs were removed, and wells were stained with 0.025% crystal violet (Thermo Fisher Scientific, Emeryville, CA) in 20% ethanol to visualize and quantify plaques. ZIKV bloodmeal titers were recorded as the reciprocal of the highest dilution where plaques were noted and are represented as PFU per mL blood.

### Zika virus vector competence experiments

Stock ZIKV inocula in DMEM or DMEM with no virus as a control, were mixed at 1:10 or 1:20 ratios with fresh heparinized sheep blood (HemoStat Laboratories, Dixon, CA) to achieve ZIKV titers of 10^4^ – 10^6^ PFU/mL for each experiment. Bloodmeals were presented to 200-300 female *Ae. aegypti* 3-5 days post-eclosion in cohorts of 100 per container with 2-3 containers per group, 24-hours after sugar withdrawal. Bloodmeals were presented for 60 minutes through a collagen membrane that was rubbed with an artificial human scent (BG-Sweetscent mosquito attractant, Biogents USA) and heated to 37°C in a membrane feeder (Hemotek Ltd, Blackburn, United Kingdom). Fully engorged females with blood in their abdomens visible at 10X magnification were cold-anesthetized by holding for 4 minutes at −20°C, sorted into clean plastic containers at a density of 20-30 mosquitoes per container, and held at 28°C with 80% humidity and 12 hour:12 hour light:dark cycle for 14 days, with constant access to filter-sterilized 10% sucrose. At 14 days post-bloodfeed, mosquitoes were cold-anesthetized and held immobile on ice. Legs and wings were removed before collection of expectorate for 20 minutes into capillary tubes containing PBS (23). Each capillary tube was placed in a 1.5-mL tube containing 250 μL PBS and centrifuged at 8,000 g for 1 minute to recover saliva. Legs/wings and bodies were placed into 2-mL tubes (Thermo Fisher Scientific, Emeryville, CA) containing 500 μL PBS and a 5-mm glass bead (Thermo Fisher Scientific, Emeryville, CA). Surgical tools were washed once in cavicide and twice in 70% ethanol between each dissection to minimize cross-contamination. For samples where microbial DNA from mosquito bodies were also analyzed in addition to viral RNA, the bodies were also washed twice in 70% ethanol and once in PBS prior to dissection to remove microbes on the surface of mosquitoes. Tissues were homogenized at 30 Hz for 10 minutes in a TissueLyzer (Retsch, Haan, Germany) before extracting viral RNA using a MagMax Viral RNA Extraction Kit (Thermo Fisher Scientific, Emeryville, CA), into 60 μL elution buffer following the manufacturer’s recommendations. Detection and quantification of viral RNA in mosquito tissues and saliva was determined by quantitative reverse transcription polymerase chain reaction (qRT-PCR) using a TaqMan Fast Virus 1-Step Mastermix and a ZIKV-specific primer set (ZIKV 1086F/1162c, probe: ZIKV 1107-FAM) using established methodologies (23, 55). Cycle threshold (Ct) values from the qRT-PCR were converted to RNA genome copies using standard curves established with known ZIKV RNA concentrations. Samples were assayed in technical duplicates and averaged together after conversion to RNA copies per mL. The limit of detection (LOD) was calculated from the standard curve linear regression line where the Ct value = 40; samples that did not yield a detectable Ct of less than 40 were reported at the LOD.

### 16S amplicon sequencing and bioinformatics

DNA from mosquitoes was extracted with a Quick-DNA Tissue/Insect Microprep Kit (Zymo Research, Irvine, CA, USA), using the manufacturer instructions, and eluted in 30 μL elution buffer. DNA from larval water and bloodmeals was extracted with a DNeasy Blood & Tissue Kit (Qiagen, Hilden, Germany) following manufacturer’s recommendations. DNA extracted from individual mosquitoes was PCR-amplified in either the V3-V4 (56), or solely the V4 (57) hypervariable region of the 16S rRNA gene. The presence and size of amplicons were confirmed by gel electrophoresis using a DNA ladder to identify amplicon size (GeneRuler 1kb Plus, Thermo Fisher Scientific, Emeryville, CA). Negative controls, including DNA extraction controls (extraction protocol with sterile PBS) and PCR controls (PCR reaction with molecular-grade H_2_O) were included in each library preparation. 16S amplicon libraries were prepared by addition of Nextera XT Index Kit v2 Set A adapter sequences (Illumina, San Diego, CA) which were cleaned using KAPA Pure Beads (Roche, Basel, Switzerland), quantified by Qubit dsDNA HS Assay (Thermo Fisher Scientific, Emeryville, CA), pooled to equimolar concentrations of 5nM per sample, and sequenced at the University of California, Davis, DNA Core Laboratory using Illumina MiSeq PE250.

Bacterial composition of individual mosquitoes from different water types and that exhibited different ZIKV infection status was assessed using bioinformatic analysis of the 16S rRNA amplicon. Paired end reads were filtered, trimmed, and processed using the DADA2 pipeline (package version 1.16.0) following the recommended workflow (58, 59), which was handed to *phyloseq* (ver 1.32.0) (60). Sequences were grouped into amplicon sequence variants (ASVs), a proxy for species (61), and assigned taxonomy using the reference database Silva v132 (62). Assigned taxa were filtered to remove environmental contaminants and sequencing artifacts. Contaminant and artifact ASVs were identified and removed if sequences were also present in the negative controls (DNA extracted nuclease-free H_2_O) or if reads aligned with “arthropod”, mitochondria, or chloroplast sequences.

Microbial ecology analyses were conducted using the R packages *phyloseq* (ver 1.32.0) and *vegan* (ver 2.5.7) (60, 63). To determine whether ASVs showed differential abundance across samples, Differential Expression Analysis was conducted using *DESeq2* (64). Random forest modelling was used to predict ASVs that distinguish mosquito cohorts, using the package *randomForest* (65). Sample reads were scaled to an even depth (mean number of reads per sample) prior to all analyses.

### Microbial quantification

Both culture-dependent and culture-independent assays were conducted in parallel to quantify live and total bacterial loads in mosquitoes and their rearing water. Culture-dependent quantification of microbes was performed by culturing 40μL rearing water or 40μL of 10-fold serial dilutions from individual mosquitoes homogenized in 500μL PBS on LB plates at 37°C for 5 days. Plated dilutions that yielded distinct, countable colonies were enumerated for each mosquito sample. Each sample was plated in technical triplicates and the mean colony count is reported. Culture-independent quantification of bacteria was performed by SYBR Green Real-Time PCR (Thermo Fisher Scientific, Emeryville, CA) to amplify the 16S rRNA gene in samples from mosquitoes and water. The mosquito data was normalized to an *Ae. aegypti* reference ribosomal protein S17 gene (RPS17) (66).

### Statistical analyses

Differences in pupation kinetics were determined by mixed effect ANOVAs with repeated measures. Bacterial abundance differences between groups were determined by either Mann-Whitney or Kruskal-Wallis tests. For 16S amplicon sequencing, differences in microbial communities were assessed using Principal Coordinate Analysis (PCoA) of weighted Unifrac distances and tested for significance with Permutational Multivariate Analysis of Variance (PERMANOVA). Quantification of the contribution of each variable to differences in microbial communities was conducted using Constrained Analyses of Principal Coordinates with the same Unifrac distances as in the PCoA analyses.

Vector competence was assessed by quantifying infection, dissemination, and transmission rates, calculated as the number of individual bodies, legs/wings, or expectorates, respectively, that yielded at detectable ZIKV RNA and then divided by the total number of individuals that ingested blood. The magnitude of ZIKV RNA in individual mosquito tissues is also reported. Differences in infection, dissemination, and transmission rates between mosquito groups were determined using Fisher’s exact tests, and differences in RNA levels were assessed by Mann-Whitney tests. Calculation of the infectious dose 50 (ID50) was performed using the nonlinear regression dose curve for LW and EW groups. All statistical analyses were performed using GraphPad PRISM 9.0.2 (GraphPad Software, San Diego, CA).

### Accession number(s)

Raw sequencing data are available from the NCBI Sequence Read Archive under BioProject entry PRJNA750810.

## Results

### Bacterial abundance and diversity decline during *Ae. aegypti* larval development

We began by assessing bacteria that persist through *Ae. aegypti* life stages. Persistence was defined as a bacterial taxon detected in more than one life stage, starting at the larval stage. A total of 31 mosquitoes reared in environmental cemetery water representing 4^th^ instar larvae (L4, n=8), pupae (n=8), and adults (1-3 days post-eclosion [dpe], n=7; >7 dpe, n=8) or pools of 100-200 eggs were sampled, and the number of bacterial ASVs was compared between individuals and to the rearing water (**Figure 1A**). Adult mosquitoes were divided into two age classes, 1-3 dpe and >7 dpe, to compare young and old adults. Bacteria were scarce in washed eggs, but significantly increased in L4 larvae (p=0.0008, Kruskal-Wallis test). Although bacterial abundance decreased across the totality of mosquito development (p=0.0002, Kruskal-Wallis), no difference in bacterial abundance between pupae and newly emerged adult females 1-3 dpe (p>0.999, Kruskal-Wallis) was detected, nor between young and old adult females >7 dpe (p=0.7802, Kruskal-Wallis). Bacterial abundance in L4 larvae was significantly lower than in adult females 7 dpe, where a decrease in mean 16S:RPS17 ratio of 129 (geometric mean=50, geometric SD=6) to 1.4 (geometric mean=0.4, geometric SD=7) (p=0.0008, Kruskal-Wallis) was detected. A total of 200 ASVs were identified across all life stages (**Table 1**), with 102 observed in water, 124 in larvae, 125 in pupae, and 99 in adults (**Supplemental Figure 1A**). Thirty-one ASVs representing 19 bacterial genera were shared between the rearing water, larvae, pupae, and adults, and most belonged to the phylum *Bacteroidetes* (**Supplemental Figure 1A-B**). The microbial community compositions across life stages were also unique, shown by distinct clustering of samples by life stage (**Figure 1B**). The microbial composition of larvae clustered close to water, while pupal compositions were more similar to those in adult mosquitoes. Concordant with the decline in microbial abundance and compositional shifts with life stage, decline in alpha diversity (total observed species and Shannon diversity indices) was also detected, with the greatest difference in in alpha diversity between L4 larvae and adults >7 dpe ([observed species] p=0.0009, [Shannon] p=0.0036, Kruskal-Wallis) (**Figure 1C**). The 10 most abundant ASVs accounted for nearly 80% of the L4 larval bacteria, with the proportion increasing to 90% as adults >7 dpe (**Figure 1D**). While *Flavobacterium* constituted the most common bacterial ASV in the rearing water (36%), *Elizabethkingia* was most common in larvae (two distinct ASVs, totaling 46%), while *Methylobacterium* expanded from 41% during pupation to 77% as adults >7 dpe. At the phylum level, *Proteobacteria* were progressively significantly enriched with each developmental stage (larva: 16±7%, pupa: 47±9%, adult 1-3 dpe: 73±9%, adult >7 dpe: 87±13%, p<0.0001, Kruskal-Wallis), such that they comprised the majority of bacteria in adult mosquitoes, despite comprising a smaller relative fraction in the rearing water (10±2%) (**Figure 1E**). Taken together, these data show that the microbes in rearing water that colonize *Ae. aegypti* are in highest abundance and diversity at the larval stage, and then decrease in relative abundance during development. A fraction of the microbes, primarily in the phyla *Proteobacteria* and *Bacteroidetes*, detected in rearing water persist and are enriched in adult *Ae. aegypti*.

**Figure 1.**
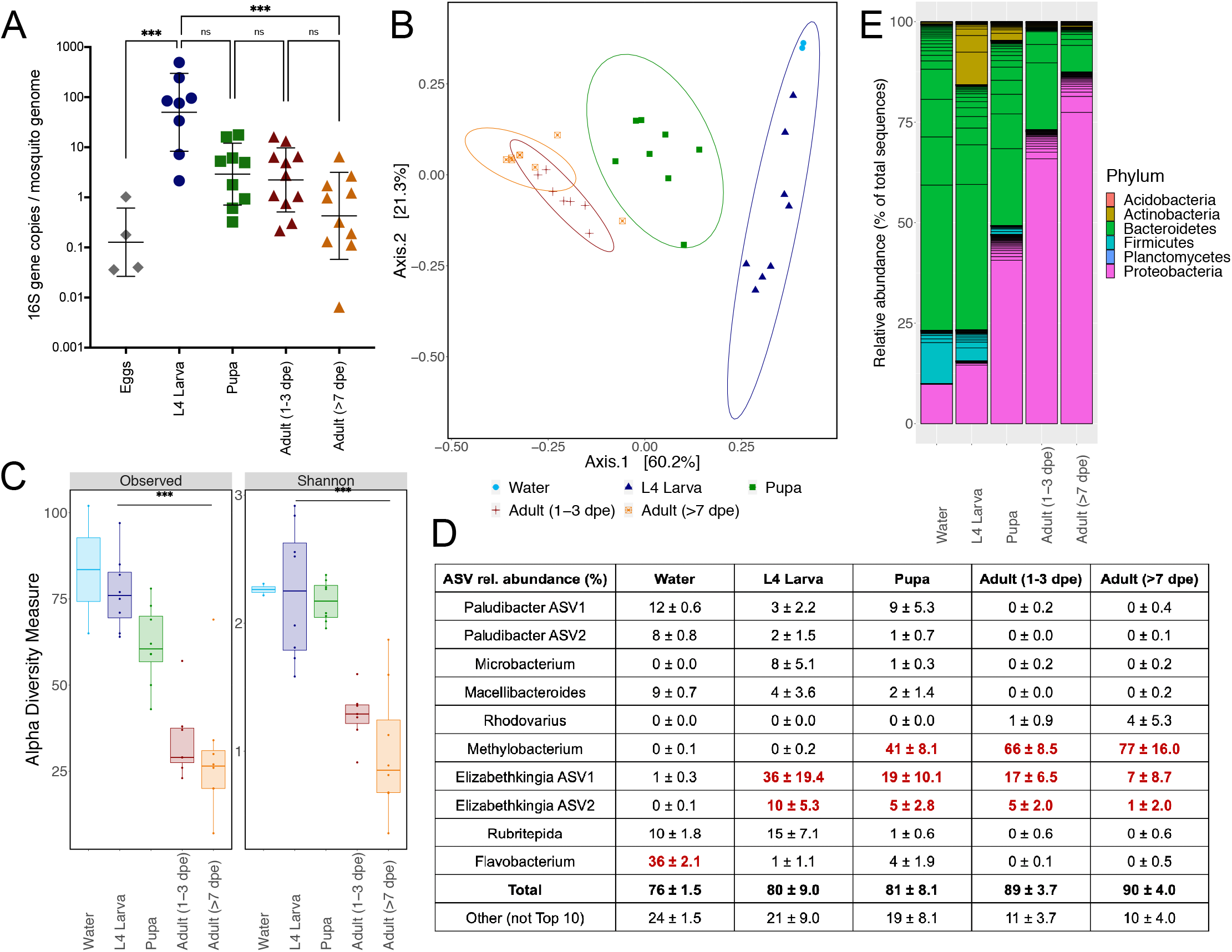
Microbial abundance decreases over the life of a mosquito although some bacterial taxa persist. A) Quantification of bacteria at each life stage of *Ae. aegypti from* Los Angeles, CA, normalized by *Ae. aegypti* gene RPS17. Each dot represents a single mosquito or a pool of 100-200 eggs. Dpe = days posteclosion. B) PCoA at the amplicon sequence variant (ASV) level, by Unifrac distances of microbial composition within each life stage. C) Alpha diversity of each life stage, using two metrics: Observed ASVs and Shannon diversity. D) Top 10 most abundant ASVs across all samples. Total indicates the sum of the top 10 ASVs, named by genus, while Other indicates the total remaining (non-top 10) ASVs. Red text highlights the ASVs that comprise the most common sequences or ASV types in the sample. E) Relative abundances of all sequences, by ASV, and colored by phylum. Sample sizes for microbiome analysis (A-E) were as follows: water (n=2, sampled at larval rearing midpoint [1 week]), mosquito (n=8 per life stage), n=4 pools of 100-200 eggs. For A and C: *** symbol denotes significance at p<0.001, using a Kruskal-Wallis test with multiple comparisons. Ns is not significant at p>0.05.

**Table 1.**
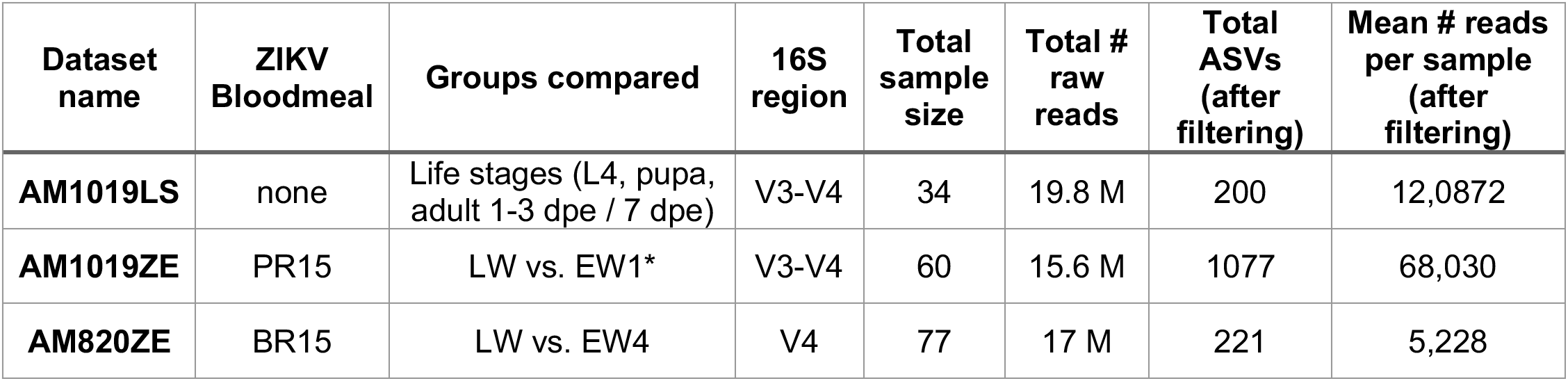
16S datasets used for microbiome analysis. All datasets included Los Angeles *Ae. aegypti* mosquitoes with bacterial DNA from their respective rearing water. *Dataset contains samples for LW, EW1, and EW2. Group EW2 was excluded as after filtering it failed to meet the threshold coverage level of 1000 reads.

### *Ae. aegypti* larvae reared in laboratory water exhibit delayed pupation relative to larvae reared in environmental water

We next asked whether the source and nature of larval water affected kinetics and success of larval development. Eggs from colonized *Ae. aegypti* were surface-sterilized, hatched, and reared to adulthood in standard laboratory water (LW) from the tap, or environment water (EW) collected outdoors from cemetery headstones (**Figure 2A**). Mosquitoes in both water types were reared at the same density and were supplemented with the same larval food quantity, which was standardized to eliminate differences in food availability and that was also sterilized to avoid introduction of additional microbes. Larvae reared in LW exhibited significantly delayed pupation, which first pupated on day 8, compared to EW-reared mosquitoes that pupated starting on day 5 (p=0.0005, paired t-test, **Figure 2B**). Furthermore, the percentage (50-81%) of larvae that pupated by day 14 in LW was significantly lower than in EW, where 100% of larvae pupated (p=0.0004, mixed effect ANOVA with multiple comparisons). LW mosquitoes pupated slower than EW mosquitoes even when the water was supplemented with baker’s yeast with or without antibiotics (mixed effect ANOVA with multiple comparisons: adjusted p=0.0023), which is conventionally used to induce hatching via hypoxia (67), and also when vacuum hatching was added, also with a goal of increasing hatch rates (mixed effect ANOVA with multiple comparisons: adjusted p=0.0022). This suggests that microorganisms in the environmental water promote pupation success and augment larval growth kinetics of *Ae. aegypti*. We also assessed whether enhanced pupation was associated with higher bacterial density in EW by comparing bacterial levels in LW to those in four EW samples (EW1-4) collected from the rearing pans 7 days post hatching. Surprisingly, bacterial DNA quantities in larval pans 7 days after hatching were not significantly different (p=0.078, Kruskal-Wallis) across LW or any EW sample (**Figure 2C**), suggesting that total microbial abundance did not influence the differences in rate of larval development to pupation. Recognizing that gene sequencing does not represent living bacteria, we also cultured and compared bacterial density in LW and EW samples, as well as in larvae, pupae, and adults (4-5 dpe) reared in both water types. The number of bacterial colonies culturable on LB agar was not significantly different between LW and EW (p=0.3143), or LW or EW reared larvae (p=0.1), pupae (p>0.99), or early adults (p=0.4286, all Mann-Whitney) (**Figure 2D**), further suggesting that the abundance of culturable bacteria does not significantly impact larval development kinetics.

**Figure 2.**
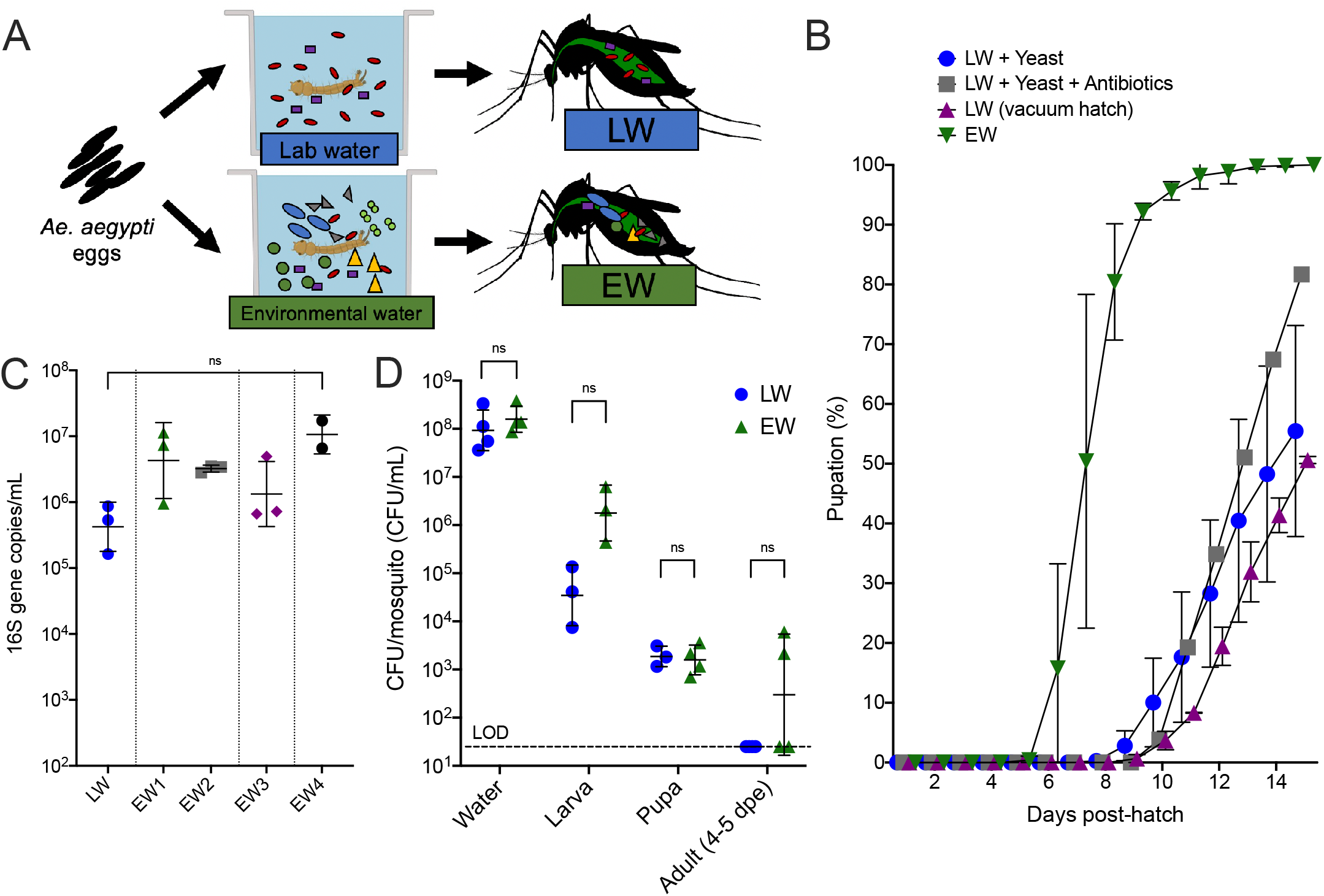
Mosquitoes reared in environmental water develop faster than those reared in laboratory water. A) Experimental design showing treatment of *Ae. aegypti* eggs with either laboratory water (LW) or environmental water (EW) from cemetery headstones. B) Pupation kinetics and rates relative to the number of larvae that hatched in cohorts of 500-600 larvae per liter. LW was spiked with either live baker’s yeast or yeast and antibiotics (penicillin-streptomycin-kanamycin, 50μg/mL). Each symbol shows the mean cumulative percent pupated on that day, with error bars denoting the range. Time-course differences in pupation were determined by mixed effects analysis (one-way ANOVA) with repeated measures and multiple comparisons. Each symbol represents the mean of replicate rearing experiments (n=3). Individual pupation rates for replicate experiments are shown in Supplemental Figure 2A. C) 16S rRNA qPCR of rearing water at 7 days posthatching. EW was collected on 4 separate occasions (EW1-4). Each symbol shows the geometric mean of PCR results from DNA extracted from 200 μL water. Values were compared by Kruskal-Wallis test with multiple comparisons. D) Colony counts of bacteria represented in colony forming units (CFU) cultured on LB agar at 37°C. Each symbol shows an average of five homogenized mosquitoes or 40μL of water at the midpoint (7 days post hatch) of a rearing experiment. The absence of colonies detected is reported at the limit of detection (LOD) of 40 CFU/mL. Pairwise comparisons between LW and EW were performed by Mann-Whitney tests.

Given that the abundance of bacteria in the larval rearing water did not explain the difference in larval growth and pupation success, we next addressed whether other differences in EW versus LW were influencing mosquito growth. To control for exogenous micronutrient content and water chemistry that could confound the observed differences in larval development, we reared larvae in diluted EW to ablate a potential pro-growth effect from EW due to these other factors. EW microbes were pelleted, washed five times in PBS, and spiked into LW at different dilutions. Although the colony-forming bacterial quantities of EW dilutions ranged from 10^2^ to 10^5^ CFU/ml at day 0 (comparable at the lowest density to 10^1.5^ CFU/ml in LW) by day 7 the bacterial numbers in all EW dilutions and LW were not significantly different (p=0.1, Kruskal-Wallis), and reached ~10^7^ CFU/mL (**Figure 3A**). Pupation rates were not different (F=5.33, p=0.07, mixed effect ANOVA) regardless of the EW dilution, and all EW groups exhibited 100% pupation by 10 dpe, which contrasted with pupation from LW where the mean was 62% (peak of 87%), which was significantly lower than all EW dilutions (F=17.19, p=0.0008, mixed effect ANOVA) (**Figure 3B**). Despite these differences in pupation rates, the quantities of colony-forming bacteria in L4 larvae were not significantly different between 1:500 or 1:10^4^ EW dilutions or LW on 7 or 10 dpe (**Figure 3C**), suggesting that larvae develop similar bacterial loads despite different initial exposure doses. The lack of difference in larval development rates at various dilutions of EW microbes, together with no difference in microbial levels across EW and LW despite augmented pupation in EW, supports specific microbes, rather than absolute microbial levels, water chemistry, or nutrient content, as a driver of faster and more efficient development of mosquitoes reared in water from the environment compared to water from the laboratory.

**Figure 3.**
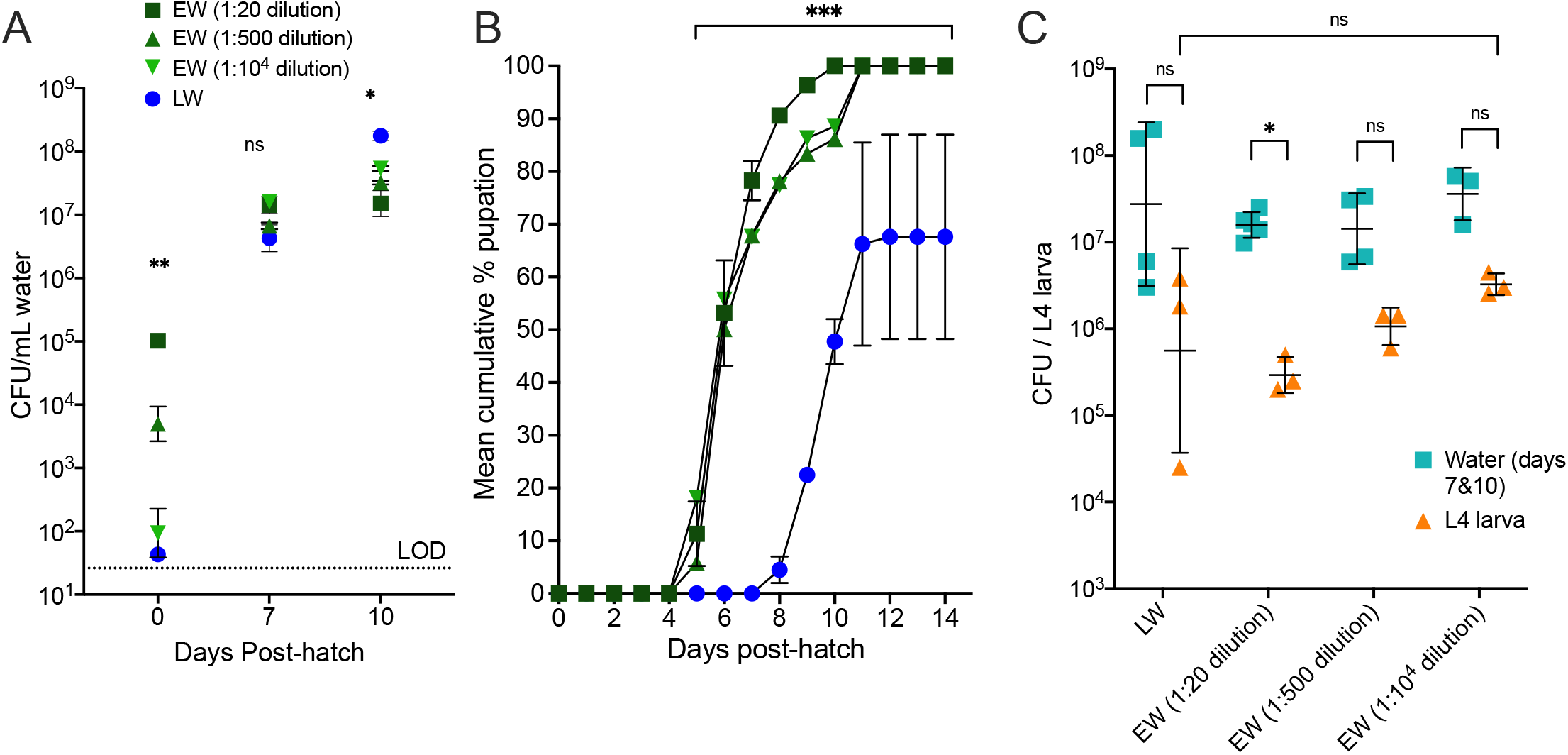
Dilution of microbes pelleted from environmental water does not delay *Ae. aegypti* larval development, which is still faster than for larvae reared in laboratory water. Microbes pelleted and washed from 3L EW were stored in glycerol stocks and diluted 1:20, 1:500, and 1:10^4^ then spiked into LW. Bacterial growth for each water treatment A) and pupation rates B) were determined. Each symbol in B shows the mean cumulative % pupation over time (individual rates for replicate experiments are shown in Supplemental Figure 2B) with error bars denoting the range. Each symbol for A shows the geometric mean of triplicate measurements and error bars denote the geometric standard deviation. Statistical tests were performed using a mixed effects analysis (one-way ANOVA) with repeated measures and multiple comparisons. C) Bacterial counts from water at days 7 and 10 were aggregated together and plotted with their respective 4^th^ instar larvae (L4) that were also sampled at the same time. Pairwise comparisons were performed using the Mann-Whitney test.

### Mosquitoes reared in environment water are less competent ZIKV vectors than mosquitoes reared in water from the laboratory

We next assessed the influence of the source of rearing water on vector competence of *Ae. aegypti* for ZIKV. LW- and EW-water reared female adult mosquitoes were presented with matched ZIKV titers or blood only in artificial bloodmeals and then assayed 14 days post-bloodfeed using qRT-PCR to detect ZIKV RNA in bodies as a marker of infection, legs and wings to indicate dissemination, and saliva to assess transmission (**Figure 4A**). No ZIKV RNA was in any mosquito that ingested blood only (data not shown). LW-reared mosquitoes were significantly more susceptible to infection and transmitted ZIKV at significantly higher rates than EW-reared mosquitoes (**Figure 4B**). This pattern was observed with 2015 ZIKV strains from Puerto Rico and Brazil and two Californian *Ae. aegypti* lineages. Although infection, dissemination, and transmission rates were higher in LW-reared mosquitoes, mean ZIKV genome copies in bodies, legs/wings, and saliva did not significantly differ between LW- and EW-reared groups (**Figure 4C** and **Supplemental Figure 3A-B** [showing additional experimental replicates that also revealed the same patterns]). Mosquitoes that contained >10^7^ ZIKV RNA copies in their body were more likely to contain detectable ZIKV RNA in saliva (LW: likelihood ratio [LR]=3.48, EW: LR=3.50) (**Figure 4D**).

**Figure 4.**
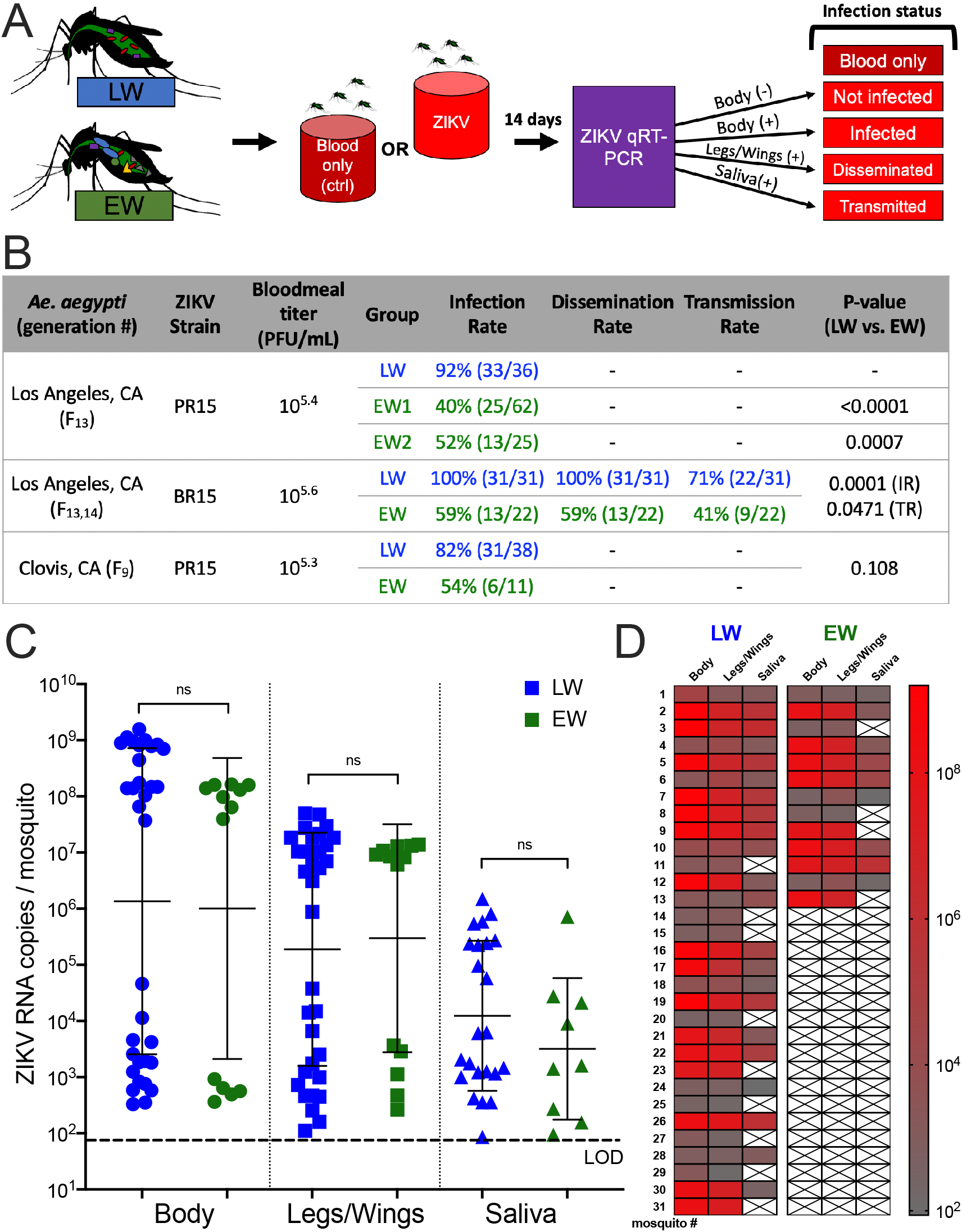
*Ae. aegypti* reared in environmental water are less competent ZIKV vectors than those reared in laboratory water. A) Experimental overview of vector competence experiments showing control (no ZIKV) and ZIKV-exposed mosquitoes reared in either laboratory water (LW) or environmental water (EW) that were incubated for 14 days post-bloodfeed and then harvested to assess infection (bodies), dissemination (legs/wings) and transmission (saliva). B) Summary table of infection, dissemination and transmission rates. Transmission rate refers to the number of individuals transmitting from the total number of individuals that ingested a bloodmeal with ZIKV. P-values were calculated with Fisher’s exact tests. C) ZIKV RNA levels in Los Angeles *Ae. aegypti* infected with ZIKV BR15. Each symbol shows a single mosquito, and only mosquitoes that were ZIKV-positive by RT-qPCR (Ct<40) are shown. Error bars denote the geometric mean and standard deviation among positive individuals. The Mann-Whitney test was used. The dotted line denotes the average limit of detection, 65 ZIKV RNA copies/mosquito or saliva sample across all RT-qPCR plates. D) Heatmap matching individual mosquitoes with their respective tissues, colored by ZIKV RNA levels.

To understand the dose response to ZIKV infection, *Ae. aegypti* reared in both water types were exposed to a range of bloodmeal titers below and above 10^5^ PFU/ml (**Figure 5A**). LW-reared mosquitoes became infected at a significantly lower bloodmeal titer than EW-reared mosquitoes (F=878, p<0.0001 for comparison of fits [slope and y-intercept], nonlinear regression) (**Figure 5B**). The infectious bloodmeal titer that produced ZIKV infections in 50% of the cohort (ID50) for LW-reared mosquitoes was 10^3.0^ PFU/ml compared to 10^5^’^6^ PFU/ml for EW-reared mosquitoes, which represents a 400-fold difference. Mosquitoes reared in both water types followed a strong dose response to ZIKV infection (R^2^ = 0.33 for LW and 0.85 for EW, nonlinear regression). Together, this data demonstrates that laboratory water-reared mosquito colonies are more susceptible to ZIKV infection and transmission than mosquitoes reared in water from the environment. The higher ID50 of EW mosquitoes also suggests that these mosquitoes are less susceptible to infection and transmission of ZIKV when ingesting a bloodmeal titer reflective of a typical human viremia (68).

**Figure 5.**
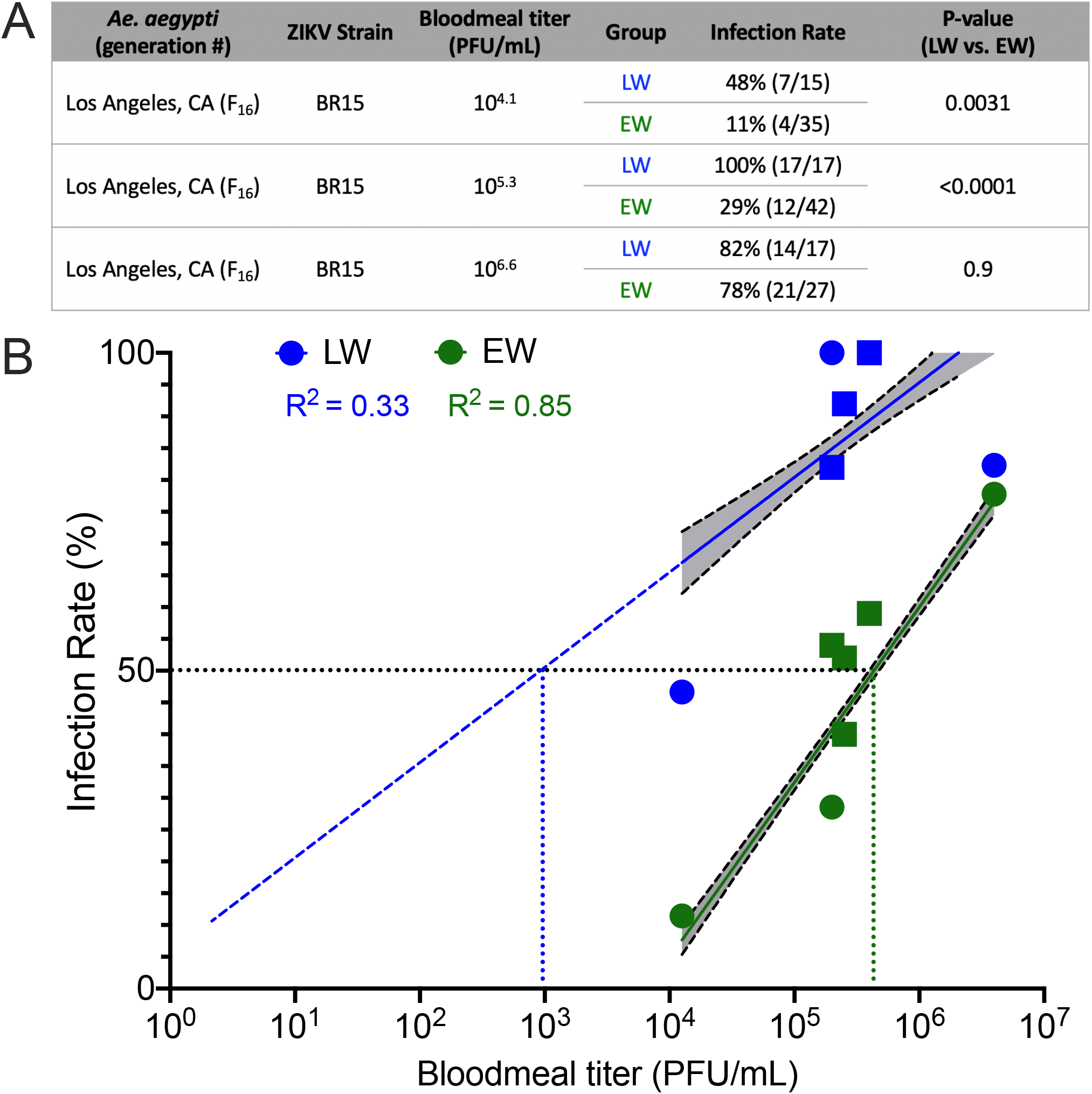
*Ae. aegypti* reared in environmental water require higher doses of ZIKV to become infected compared to those reared in laboratory water. A) Summary of additional vector competence experiments using Los Angeles *Ae. aegypti* and ZIKV BR15 at three different bloodmeal titers. B) Plot of infection rate versus bloodmeal titer. Squares represent the same data in Figure 4B and circles indicate the additional experiments in Figure 5A. A best-fit non-linear regression line, with 95% confidence intervals is shaded in grey. LW: slope = 14.9 [95%CI = 11.5–18.3], y-intercept = 5.949 [95%CI = −12.57–24.44]; EW: slope = 27.6 [95%CI = 26.01–29.01], y-intercept = −105.3 [95%CI = −113.8– −96.83].

### Larval water source does not differentiate bacterial composition between adult mosquitoes as much as bloodmeal status

Although LW and EW *Ae. aegypti* that were not ZIKV-exposed showed similar bacterial levels, we next questioned whether the same pattern would be observed in the context of ZIKV infection. Adult female mosquitoes reared in LW or EW that ingested ZIKV in bloodmeals were grouped into the following classes based on their infection outcomes: 1) not infected, where no ZIKV RNA was detected above the limit of detection of 65 ZIKV genomes/body, 2) infected (low), defined as body titers <10^6^ ZIKV genomes/body, or 3), infected (high), defined as body titers >10^6^ ZIKV genomes/body. The ‘high’ and ‘low’ infection states were defined based on the bimodal distribution of RNA levels observed in bodies (**Figure 4C**). The reasoning for this grouping is that individuals with high ZIKV RNA levels in their bodies were more likely to have disseminated infections that lead to ZIKV RNA detection in saliva (**Figure 4D**), a pattern also observed in previous studies with *Ae. aegypti* from the same source colonies and that used the same ZIKV strains as in this study (23). Mosquitoes that fed on blood without ZIKV or that had only been presented with sugar water were included as controls. Prior to a bloodmeal, where mosquitoes had been exposed to only sugar at 3 dpe, both LW and EW females had bacterial quantities in their bodies that were not significantly different (p=0.5476, Mann-Whitney) and the bacterial load was low (**Figure 6A**). Ingestion of blood resulted in a 50-100-fold increase in bacterial levels in both groups compared to their unfed mosquitoes of the same age (p=0.0005, Kruskal-Wallis [multiple comparisons: LW – adjusted p=0.012; EW – adjusted p=0.0212]). Bloodfed LW mosquitoes contained significantly higher bacterial levels compared to EW mosquitoes (p = 0.0079, Mann-Whitney). Regardless of infection outcome, both LW and EW mosquitoes that ingested ZIKV showed lower bacterial levels than blood-only groups (F=33.41, p<0.0001, F=40.30, p<0.0001 for infection state and water type, respectively, two-way ANOVA). Moreover, the two-way ANOVA on bloodfed mosquitoes detected a significant interaction between water type and ZIKV infection state (interaction F=4.6, p=0.0032). LW mosquitoes that were not ZIKV infected or that were infected at high levels contained higher bacterial levels compared to EW mosquitoes (p=0.0043, p=0.0095, respectively, Mann-Whitney).

**Figure 6.**
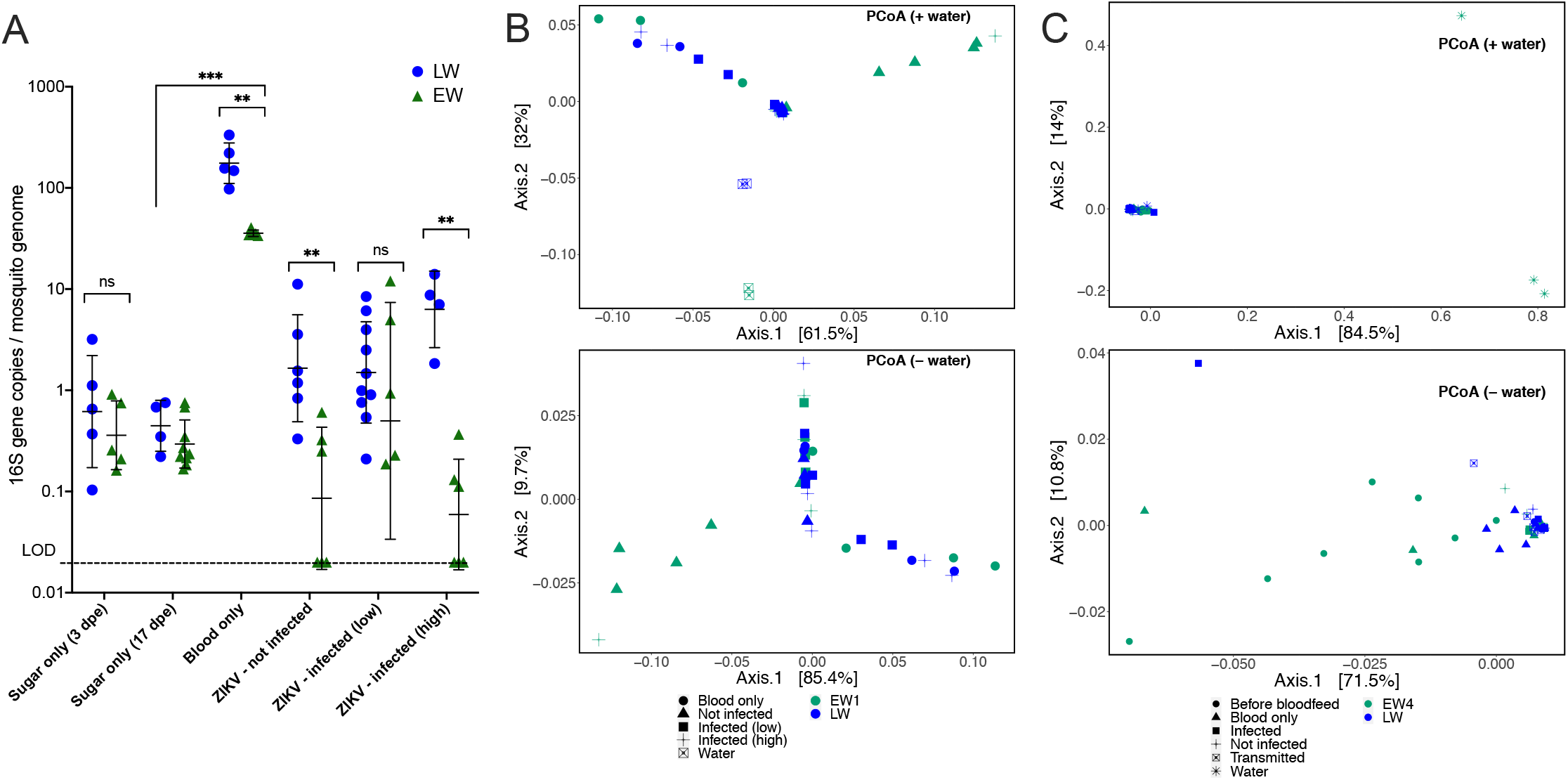
Microbial communities differ slightly by water type. A) Bacteria in LW- or EW-reared *Ae. aegypti* exposed to sugar, blood only, or blood containing ZIKV quantified by 16S qPCR and normalized to *Ae. aegypti* reference gene RPS17. Symbols refer to a single mosquito, and error bars denote the geometric mean and geometric standard deviation. Statistically significant differences between LW and EW were determined by Mann-Whitney test, while differences across treatments were determined by Kruskal-Wallis test. Interaction effects between water type (LW, EW) and ZIKV infection states were investigated by two-way ANOVA with multiple comparisons, on logio-transformed values. (*p<0.05, **p<0.01, ***p<0.001, ns is not significantly different at p=0.05 level). B) and C) depict the Principal Coordinates of Unifrac distances, colored by water type. B shows a cohort of Los Angeles *Ae. aegypti* presented with ZIKV PR15 (dataset AM1019ZE, Table 1) while C shows another cohort of Los Angeles *Ae. aegypti* presented with ZIKV BR15 (dataset AM820ZE, Table 1). Top plots include mosquito and rearing water samples while bottom plots have water samples omitted for higher resolution.

Next, we examined the bacterial composition of LW and EW mosquitoes, reasoning that the type of bacteria may influence vector competence more than the total bacterial load. We compared the relative bacterial abundances and taxonomic diversity of LW and EW adult mosquitoes as well as across mosquitoes that exhibited differential ZIKV infection states from two replicate vector competence experiments. A total of 1077 ASVs and 221 ASVs were identified in the first and second vector competence experiments, respectively (AM1019ZE and AM820ZE, **Table 1**), most of which were detected in the rearing water (**Supplemental Figure 4**). Comparing EW1 and LW, we detected no significant differences in clustering of samples by water type; however, when we compared EW4 and LW, we detected a difference in clustering (PERMANOVA: [F=2.52, p=0.043, Adonis], [F=6.91, p=0.018, Betadisper]). The disparity in clustering patterns across different EW sample collections may be due to differences in community distribution in EW4 vs. LW arising from the low number of ASVs, where most EW4 and LW mosquitoes in the AM820ZE dataset were dominated by *Psuedomonas* (**Figure 6C**, **Supplemental Figure 4C, 6C**). Despite the identification of different ASVs between experimental iterations, EW had higher bacterial diversity and evenness than LW ([richness: F=17.25, p<0.0001; Shannon: F=12.8, p<0.0001], one-way ANOVA) (**Supplemental Figure 4**). When we only compared mosquitoes (and not their rearing water), the bacterial compositions between LW and EW mosquitoes were slightly different, as shown with partial sample overlaps, although clustering was not significantly different (PERMANOVA: [F=1.87, p=0.111, Adonis], [F=0.15, p=0.707, Betadisper]) (**Figure 6B**, bottom panel). Several EW mosquitoes that were refractory to ZIKV infection clustered together; these individuals had an increased proportion of *Asaia* and *Flavobacterium* and a significantly reduced proportion of *Rhodovarius, Micrococcus*, and *Neochlamydia* bacteria compared to infection competent individuals, as determined by DESeq2 analysis and random forest modelling (**Supplemental Figure 5A, 6A-B**). The overall contribution of water type to differences in bacterial composition across individual mosquitoes of either bloodfed status was 22%. Whether mosquitoes ingested ZIKV and whether mosquitoes that ingested ZIKV became infected were also important variables that explained 27% and 34%, respectively, of the differences in bacterial composition across groups (**Table 2**). EW mosquitoes that ingested blood with ZIKV had a reduced abundance of *Serratia* compared to EW mosquitoes that ingested blood only (**Supplemental Figure 5B, 6B**). For both datasets, there were no significant differences in bacterial compositions in LW mosquitoes that ingested blood only or ZIKV; this may be an artifact of LW mosquitoes possessing few bacterial taxa such that differential abundances could not be detected (**Supplemental Figure 6A, B, Table 2**). Additionally, no differences in bacterial composition were detected between mosquitoes with high and low levels of ZIKV RNA or between mosquitoes that were infected and those that transmitted (**Figure 6C, Table 2**). Taken together, EW-reared adult females harbor different microbiota when reared in the same water source collected at different times. This suggests that despite differences in microbiota, adult female *Ae. aegypti* exhibit a consistently reduced vector competence for ZIKV when reared in environmental water compared to laboratory water.

**Table 2.** Summary of contributions to microbial compositional differences by mosquito variable. Variable contribution, the percentage of variance between samples associated with the metadata, was calculated using constrained analysis of Principal Coordinates. Statistical tests were calculated by permutational ANOVA (p<0.05, p<0.01).

| Group | Variable compared 1 | Variable compared 2 | Variable contribution | p-value |
| --- | --- | --- | --- | --- |
| - | EW1 | LW | 22% | 0.008** |
| EW1 | ZIKV-exposed | Blood only | 27% | 0.019* |
|  | Infected | Not infected | 34% | 0.011* |
|  | Infected (low) | Infected (high) | 10% | 0.676 |
| LW | ZIKV-exposed | Blood only | 13% | 0.619 |
|  | Infected | Not infected | 8% | 0.438 |
|  | Infected (low) | Infected (high) | 10% | 0.676 |
| - | EW4 | LW | 5% | 0.012* |
| EW4 | Before bloodfeed | After bloodfeed | 11% | 0.074 |
|  | ZIKV-exposed | Blood only | 12% | 0.01** |
|  | Infected | Not infected | 8% | 0.463 |
|  | Infected | Transmitted | 9% | 0.963 |
| LW | Before bloodfeed | After bloodfeed | 6% | 0.692 |
|  | ZIKV-exposed | Blood only | 9% | 0.037* |
|  | Infected | Not infected | 7% | 0.319 |
|  | Infected | Transmitted | 5% | 0.693 |

## Discussion

Here we show that microbial diversity stemming from different water sources used to rear larvae in a laboratory environment modifies vector competence of *Ae. aegypti* for ZIKV. Reduced vector competence in environmental water-reared *Ae. aegypti* was consistently observed using two lineages of Californian *Ae. aegypti* and two epidemiologically relevant ZIKV strains. These results suggest that modification of *Ae. aegypti* developmental conditions to reflect environmental water as compared to laboratory tap water, which is conventional, decreases laboratory infection and transmission rates. Use of laboratory water to rear larvae likely leads to overestimates of the transmission potential of ZIKV vectors in the environment. This pattern may apply to other vector-virus pairings as well; future research should address this question.

Differences in pupation kinetics between EW-versus LW-reared mosquitoes indicate that the type of bacteria, but not bacterial abundance, impacts success of mosquito larval development; this mostly agrees with previous studies on gnotobiotically-reared larvae with bacteria and yeast of similar densities (69). Our observation of wide variability in microbial content in experiments using different collections of environmental water, but which all yielded 100% pupation success, suggests that there is likely functional redundancy in microbes needed to nutritionally support larval growth and stimulate pupation. While the bacterial diversity of laboratory and environmental water from natural mosquito larval habitats were different, bacterial taxonomic differences within mosquitoes reared in water from these respective sources were more subtle. This suggests that mosquitoes may harbor a relatively low number of species in a “core” microbiome (70), possibly explaining the sparse number of bacterial species detected and the lack of shared species across experimental replicates. In concordance with previous *Ae. aegypti* microbiome studies, we observed a high relative abundance of *Proteobacteria* and *Bacteroidetes* in adult mosquitoes (38, 42). At the genus, level, most adult mosquitoes were dominated by *Asaia, Flavobacterium, Elizabethkingia*, and *Pseudomonas* bacteria. These bacteria were also found in small quantities in their rearing water, suggesting they are likely environmental in origin, except for *Elizabethkingia*, which was also detected in surface-sterilized eggs. Because the same ASVs matching to *Elizabethkingia* were also identified in surface-sterilized eggs, the origin of *Elizabethkingia* in mosquitoes in this study cannot be determined. Bacteria from this genus are present in the environment, larvae, newly emerged adults, and also in reproductive tissues of *Aedes spp*. mosquitoes (71), but have not yet been reported in eggs.

By varying the source of larval rearing water, we aimed to modify the microbiota of *Ae. aegypti* with the premise that mosquito microbes are acquired through the environment and especially larval water. We therefore expected that a sterile sugar diet and a single artificial bloodmeal provided to adults would narrow the microbial input of the mosquitoes to reflect larvae-acquired microbes from the rearing water. While we detected differences in the microbiota in LW-versus EW-reared mosquitoes, the microbiota was more different between control bloodfed and ZIKV-bloodfed groups. Other studies have also measured strong relationships between bloodmeal status and microbiome composition (72, 73), some showing greater differences in expression of *Ae. aegypti* genes in mosquitoes that bloodfed than between axenic and conventionally reared mosquitoes (74). A functional limitation to this and previous work is the inability to account for all microbial sources in adult mosquitoes stemming from their natural field environment, including microbes acquired during sugar feeding of adults on flora.

Despite the lack of reproducible changes in species composition of bacteria in adult *Ae. aegypti* reared in different aquatic environments, we observed a substantial effect on vector competence, where EW-reared mosquitoes exhibited lower infection and transmission rates than LW-reared mosquitoes. As this is the first study examining the microbiota of Californian *Ae. aegypti*, and one of few mosquito studies using ASVs instead of operational taxonomic units (OTUs), where ASVs are gaining favor over OTUs due to their increased taxonomic resolution as well as their consistent labelling (61), direct comparisons to other *Ae. aegypti* microbiome studies should be made with caution. In addition to a “core microbiome” effect on mosquito vector competence, there could also be functional redundancy in the effects of microbiota on mosquito physiology. Despite microbial variability in rearing water and mosquitoes observed in our experimental replicates, increased infection and transmission of ZIKV by LW-compared to EW-reared mosquitoes was reproducible.

An overall reduction of bacterial levels in ZIKV-exposed mosquitoes relative to non-exposed mosquitoes suggests that ZIKV infection negatively impacts the mosquito microbiota. This could be due to interactions between the mosquito antiviral immune response and a generalized antimicrobial effect that indirectly kills bacteria within the mosquito gut. Previous work on *Ae. aegypti* innate immunity implicated a link between antiviral and antibacterial immune responses to infection (48, 75, 76). For example, the Toll pathway, which recognizes bacterial cell walls in insects, also modulates responses to DENV infection (47). This implicates a nonspecific pan-arboviruses effect, where elevated immune responses in response to resident microbiota confer resistance to infection. Furthermore, additional life history traits like adult body size are influenced by larval water conditions (16, 50), implicating a physiological modification that may indirectly result from microbial exposures of larvae. Moreover, gut microbes play a nutritional role in mosquito symbiosis (44, 74), and larval nutrition impacts mosquito size and development (77), although the role of size in vector competence of *Ae. aegypti* and DENV and *Culex spp*. mosquitoes and West Nile virus is controversial (78–80). Carryover effects (49) of larval exposure to isolates of *Flavobacterium, Lysobacteria, Paenibacillus*, and *Enterobacteriaceae* on adult lipid metabolism and DENV infection in *Ae. aegypti* that corroborate our observations that bacterial exposure during the larval stage can influence adult mosquito traits. The influence of bacteria known to impact vector competence in a monoculture in the context of the complex microbial community should be a target of future research.

## Acknowledgements

WL acknowledges funding support from the Training Grant Program of the Pacific Southwest Regional Center of Excellence for Vector-Borne Diseases funded by the U.S. Centers for Disease Control and Prevention (Cooperative Agreement 1U01CK000516) and from the University of California, Davis, School of Veterinary Medicine Graduate Student Support Program.

---

**Supplemental Table 1.** Water sources, collection dates, and data references for *Ae. aegypti* used in rearing experiments. LW is laboratory water, and EW is environmental water.

| Water type | Collection source | Collection date | Figures (16S libraries) |
| --- | --- | --- | --- |
| EW1 | Davis, CA, USA<br>cemetery headstone | Aug 2018 | Fig 1, 6B, Supplemental 3A,<br>Supplemental 4A-B (AM1019LS,<br>AM1019ZE) |
| EW2 |  | Jun 2019 | Fig 4B, Supplemental 3A |
| EW3 |  | Nov 2019 | Supplemental Fig 3B |
| EW4 |  | Feb 2020 | Fig 6C, Supplemental 4C-D<br>(AM820ZE) |
| LW | Insectary tap<br>(deionized) | At experiment<br>start date | Matched with equivalent EW |

**Supplemental Figure 1.**
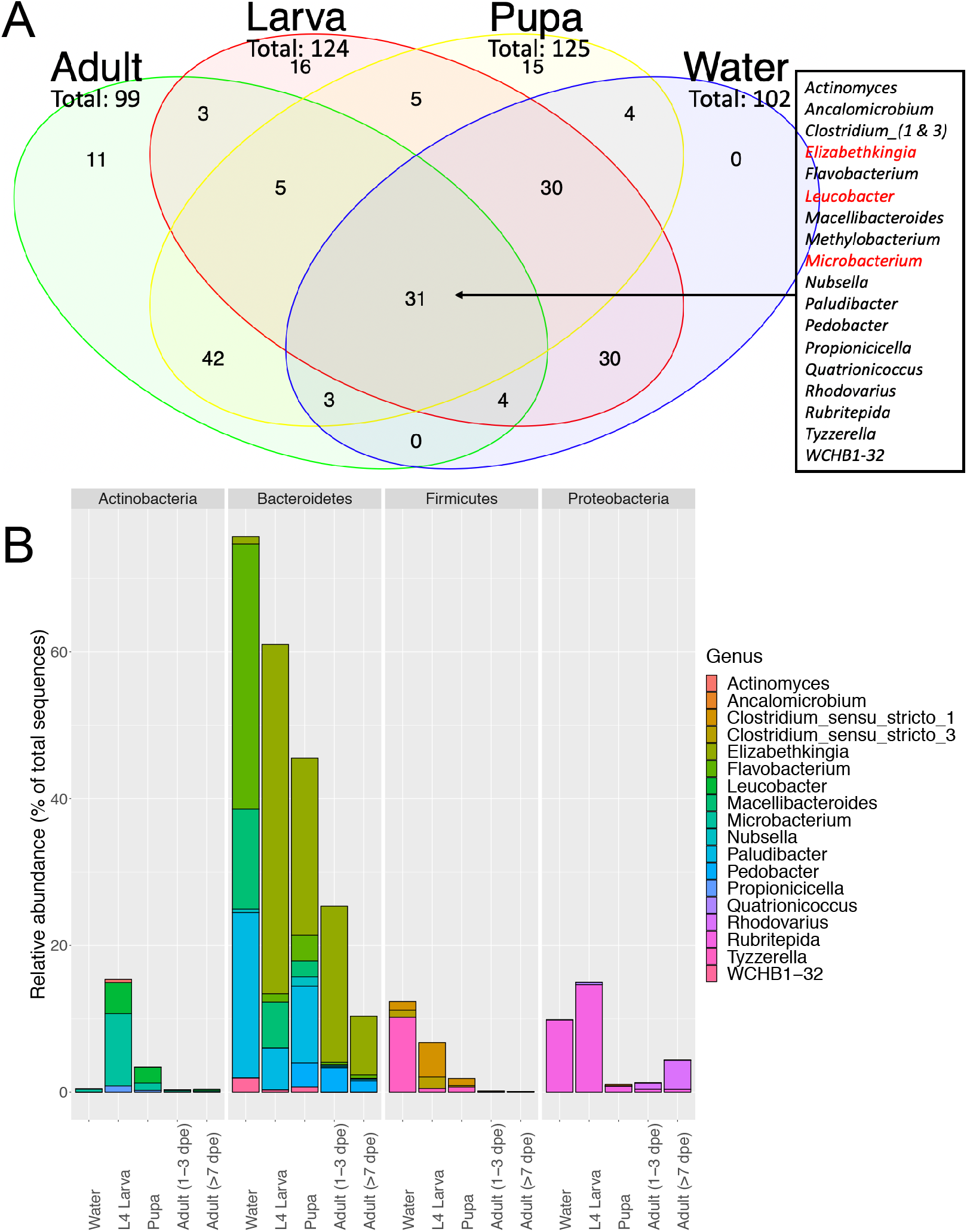
Bacterial ASVs shared in various life stages. A) Shared and distinct ASVs detected in rearing water, larvae, pupae, and adult mosquitoes. Text box lists shared bacteria by genus. Red text indicates genera that were also detected in eggs after surface sterilization. B) Relative abundance of the 19 genera shared among water, larvae, pupae, and adult mosquitoes grouped by bacteria taxa.

**Supplemental Figure 2.**
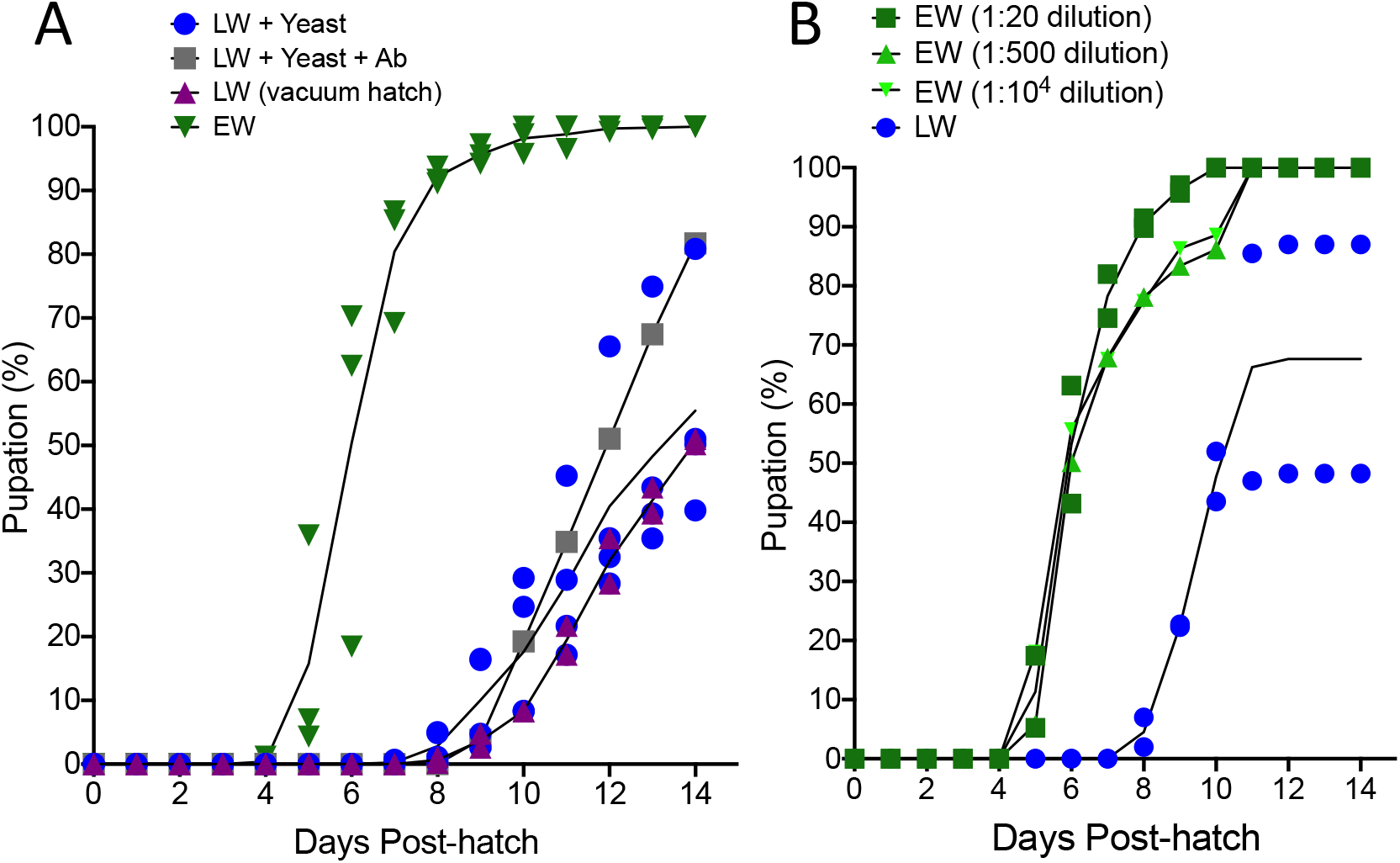
Development times of Los Angeles *Ae. aegypti*, plotted by individual experimental replicates. Pupation differences between A) EW vs. LW variations in Fig 2B, and B) LW vs. EW dilutions in Fig 3B. Black lines denote the average % pupation of replicate experiments.

**Supplemental Figure 3.**
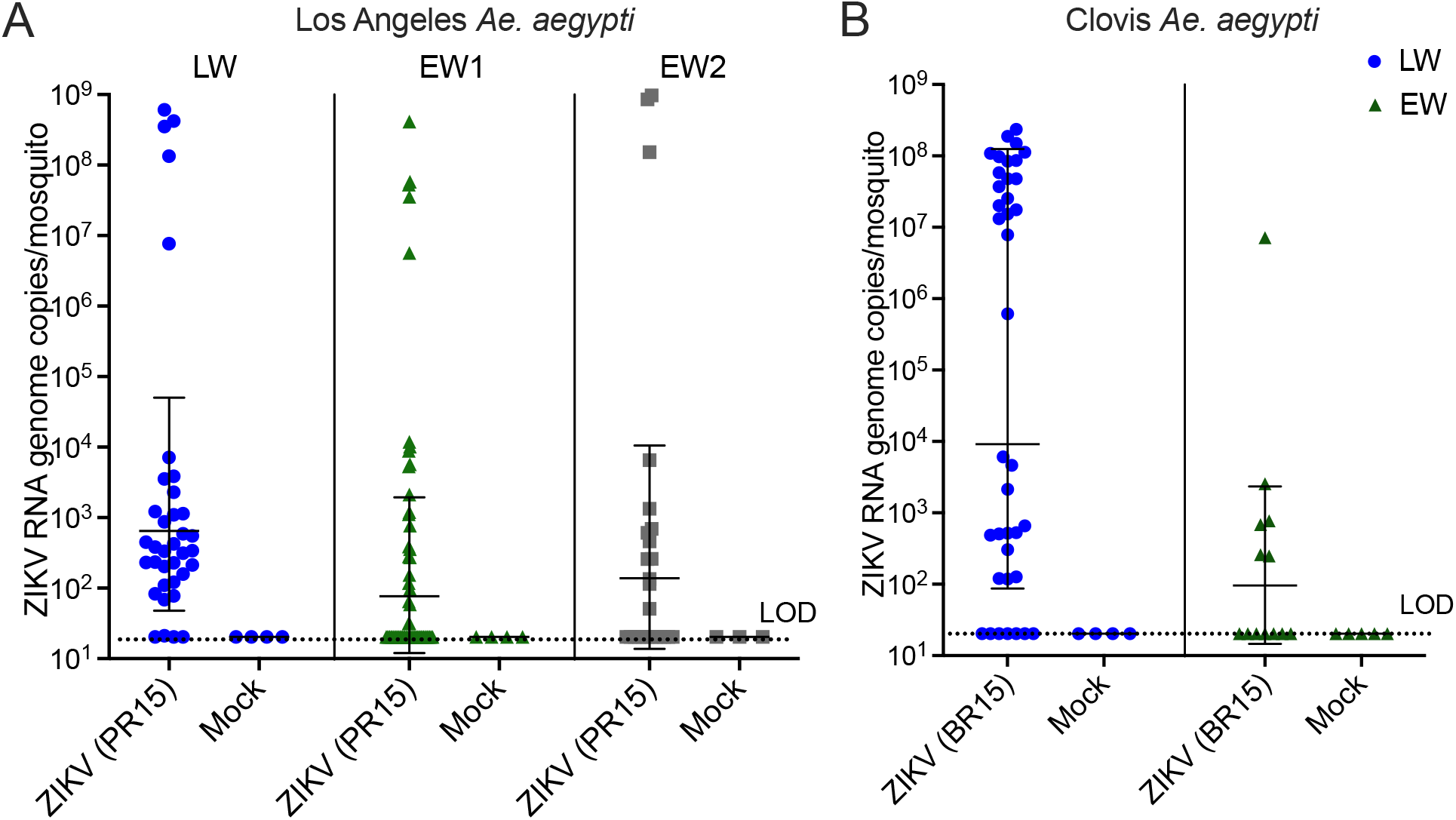
ZIKV RNA levels in individual mosquitoes that ingested bloodmeals containing 10^5.4^ (A) or 10^5.3^ (B) PFU/ml ZIKV or blood with no virus (mock), summarized in Fig 4B. A) *Ae. aegypti* colonized from Los Angeles, CA were presented with PR15 ZIKV. B) *Ae. aegypti* colonized from Clovis, CA were presented with ZIKV BR15. Mosquitoes with no detectable ZIKV RNA are reported at the limit of detection (LOD) of the assay, which averaged 10^1.3^ ZIKV genome copies/mosquito in both A and B.

**Supplemental Figure 4.**
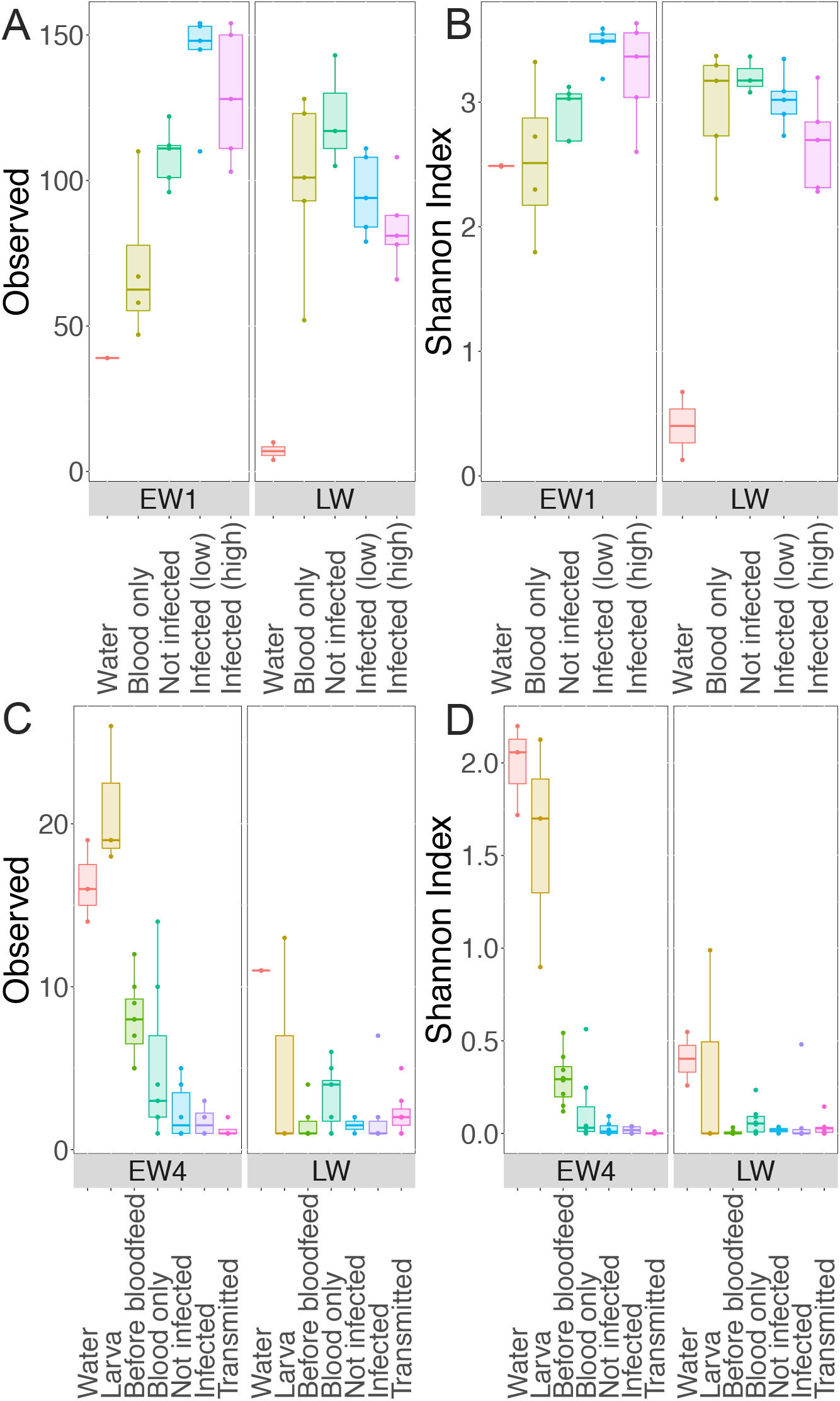
Alpha diversity of EW1, EW4, and LW mosquitoes.

**Supplemental Figure 5.**
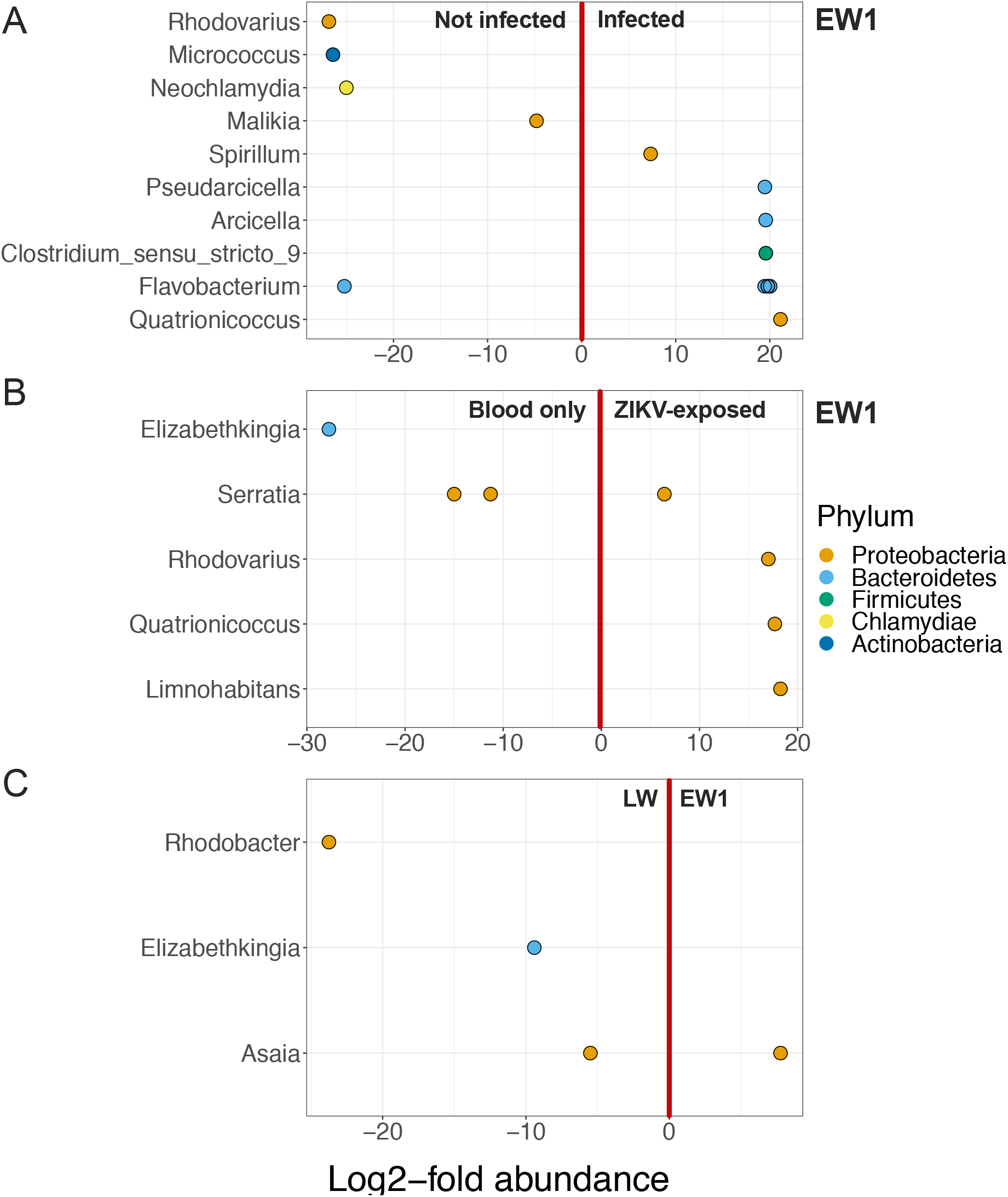
Differentially abundant taxa determined by DESeq2. Each dot represents an ASV from the respective genus (y-axis), that was differentially abundant among mosquitoes according to infection status or water type which are separated by the vertical red line. ASVs to the left of the dividing line were more abundant in the left group while ASVs to the right of the dividing line were more abundant in the right group. Bacterial taxa from *DESeq2* were significantly different if the adjusted p-value cut-off (alpha) was below 0.05. No differentially abundant ASVs were found between Not infected/ Infected and Blood-only/ZIKV-exposed mosquitoes in LW mosquitoes.

**Supplemental Figure 6.**
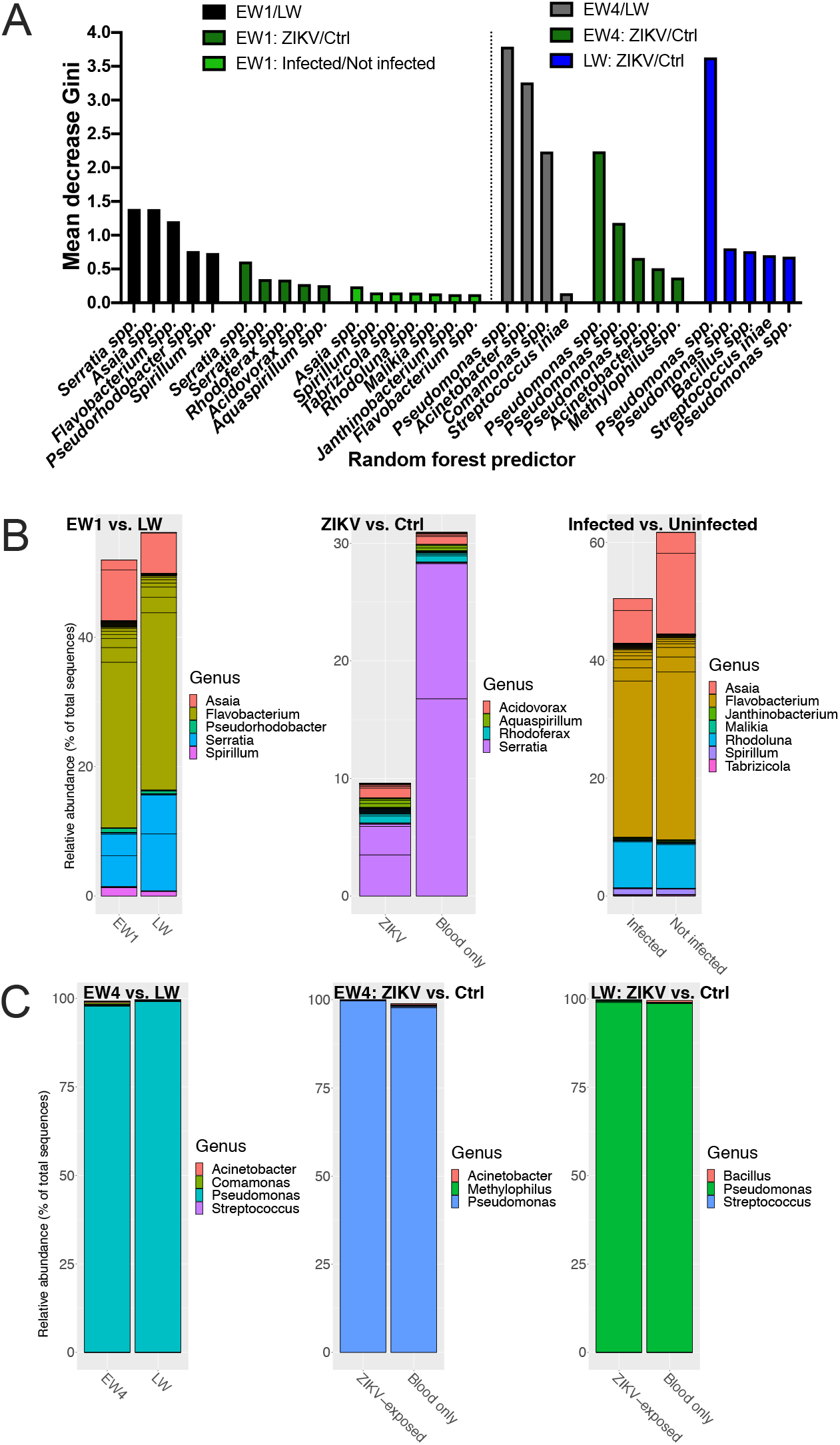
Random forest modelling of significant bacterial taxa. A) Top 5-7 predictors of mosquito groups were plotted, and B) relative abundance of the top 5-7 selected predictors were plotted using ggplot2. ‘Ctrl’ indicates mosquitoes that ingested blood only but no ZIKV.

